# Spreading predictability in complex networks

**DOI:** 10.1101/2020.01.28.922757

**Authors:** Na Zhao, Jian Wang, Yong Yu, Jun-Yan Zhao, Duan-Bing Chen

## Abstract

Spreading dynamics analysis is an important and interesting topic since it has many applications such as rumor or disease controlling, viral marketing and information recommending. Many state-of-the-art researches focus on predicting infection scale or threshold. Few researchers pay attention to the predicting of infection nodes from a snapshot. With developing of precision marketing, recommending and, controlling, how to predict infection nodes precisely from snapshot becomes a key issue in spreading dynamics analysis. In this paper, a probability based prediction model is presented so as to estimate the infection nodes from a snapshot of spreading. Experimental results on synthetic and real networks demonstrate that the model proposed could predict the infection nodes precisely in the sense of probability.

## Introduction

Spreading dynamics is an important issue in spreading and controlling [1–3] of rumor [4–7] and disease [8–11], marketing [12], recommending [13–15], source detecting [16, 17], and many other interesting topics [18–22]. How to predict the infection probability [23], infected scale [24, 25], and even the infected nodes precisely has been gotten much attention in recent years.

Researchers have gotten many achievements on macro level of spreading such as phase transition of spreading [26] and basic reproduction number [27]. Up to now, many researches focus on estimating of infection scale. The simplest one is mean-field model, in which, the spreading coverage can be predicted by using differential equations [24]. Besides mean-field model, some more realistic models such as pair approximation [25] and permutation entropy [28] are considered to predict the infection scale or infectious disease outbreaks. The main difference between mean-field and pair approximation is that the former(latter) approximates high-order moments in term of first (second) order ones. In [28], the researchers studied the predictability of a diverse collection of outbreaks and identified a fundamental entropy barrier for disease time series forecasting through adopting permutation entropy as a model independent measure of predictability. Funk et al [29] presented a stochastic semi-mechanistic model of infectious disease dynamics that was used in real time during the 2013–2016 West African Ebola epidemic to fit the simulated trajectories in the Ebola Forecasting Challenge, and to produce forecasts that were compared to following data points. Venkatramanan et al [30] proposed a data-driven agent-based model framework for forecasting the 2014–2015 Ebola epidemic in Liberia, and subsequently used during the Ebola forecasting challenge. The data-driven approach can be refined and adapted for future epidemics, and share the lessons learned over the course of the challenge. Zhang et al [31] proposed a measurement to state the efforts of users on Twitter to get their information spreading. They found that small fraction of users with special performance on participation can gain great influence, while most other users play a role as middleware during the information propagation.

Up to now, most researches are focused on macro level of spreading prediction, but few on micro level. However, the detailed infected individuals should be known so as to contain the spread of serious infectious diseases such as SARS [32, 33] and H7N7 [34, 35]. Besides aspect of macro level of spreading, we should pay attention to some more details besides the general infection coverage so as to achieve fine prediction. Chen et al. did some interesting works on this area [23]. They presented an iterative algorithm to estimate the infection probability of the spreading process and then apply it to mean-field approach to predict the spreading coverage.

Combing mean-field or pair approximation models with infection probability estimating strategy [23], the number of infected nodes from a given snapshot of the propagation on network can be predicted, but can not determine which nodes will be infected. In this paper, we present a probability based prediction model to estimate the infection probability of a node, further, to determine the nodes being infected in the future.

## Materials and methods

For a given snapshot, a susceptible node can be infected by a probability in the future. Denoting by *P*_*u*_(*t*) the score of node *u* at time *t*, we have,

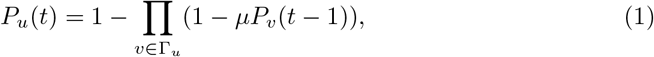

where Γ_*u*_ is the neighbors of node *u* and infected probability *µ* is estimated by IAIP model (Iterative Algorithm for estimating the Infection Probability) [23]. Since an infected node always attempts to infect its susceptible neighbor once time and a recovered node doesn’t infect any of its susceptible neighbor, so, in Eq. (1), for node *v*, it is reasonable to assume that *P*_*v*_(*t*) = 1 for infected node and *P*_*v*_(*t*) = 0 for recovered node. For susceptible node *u*, the probability to be infected at time *t* is *P*_*u*_(*t*).

Obviously, the initial condition is,

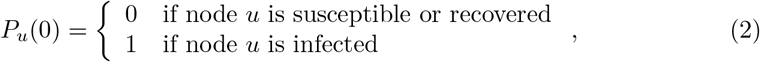

In Eq. (1), the score *P*_*u*_(*t*) for susceptible node *u* will be converged to a unique steady state denoted by *P*_*u*_(*t*_*c*_), where *t*_*c*_ is the convergence time. The final score *P*_*u*_ = *P*_*u*_(*t*_*c*_) is the probability to be infected of susceptible node while spreading achieves steady state.

Fig. 1 is a toy network with 24 nodes. The snapshot includes 5 recovered nodes and 1 infected node, as shown in Fig. 1(a). A certain spreading simulation result, average result on 10000 simulations, and result of probability prediction model from snapshot are shown in Figs. 1(b), (c) and (d) respectively. From this toy network, it can be seen that the result obtained by the probability prediction model is coincident with that by the average over 10000 simulations very well, that is, nodes 7, 8, and 19 have high probability to be infected, nodes 2 and 9 have middle probability to be infected, while other nodes have relatively low probability to be infected, as shown in Fig. 1(c) and (d). At the same time, Figs. 1(b) and (c) reflect the correlation between a certain spreading simulation and average over 10000 simulations. In order to describe how well a certain spreading simulation relative to average over 10000 simulations and result obtained by probability prediction model relative to the result of average over 10000 simulations, we use predictability *χ* and Pearson correlation *ρ* to evaluate our model. These two metrics can be calculated by:

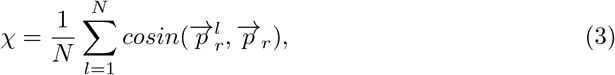

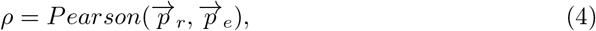

where 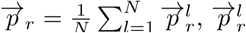 is the vector of infected frequency of nodes on the *l*^*th*^ simulation and 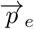 is the vector of infected probability of nodes obtained by probability prediction model. The element 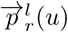 of 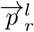 is determined by:

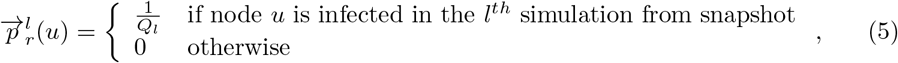

where *Q*_*l*_ is the number of infected nodes of the *l*^*th*^ simulation from snapshot.

**Fig 1.**
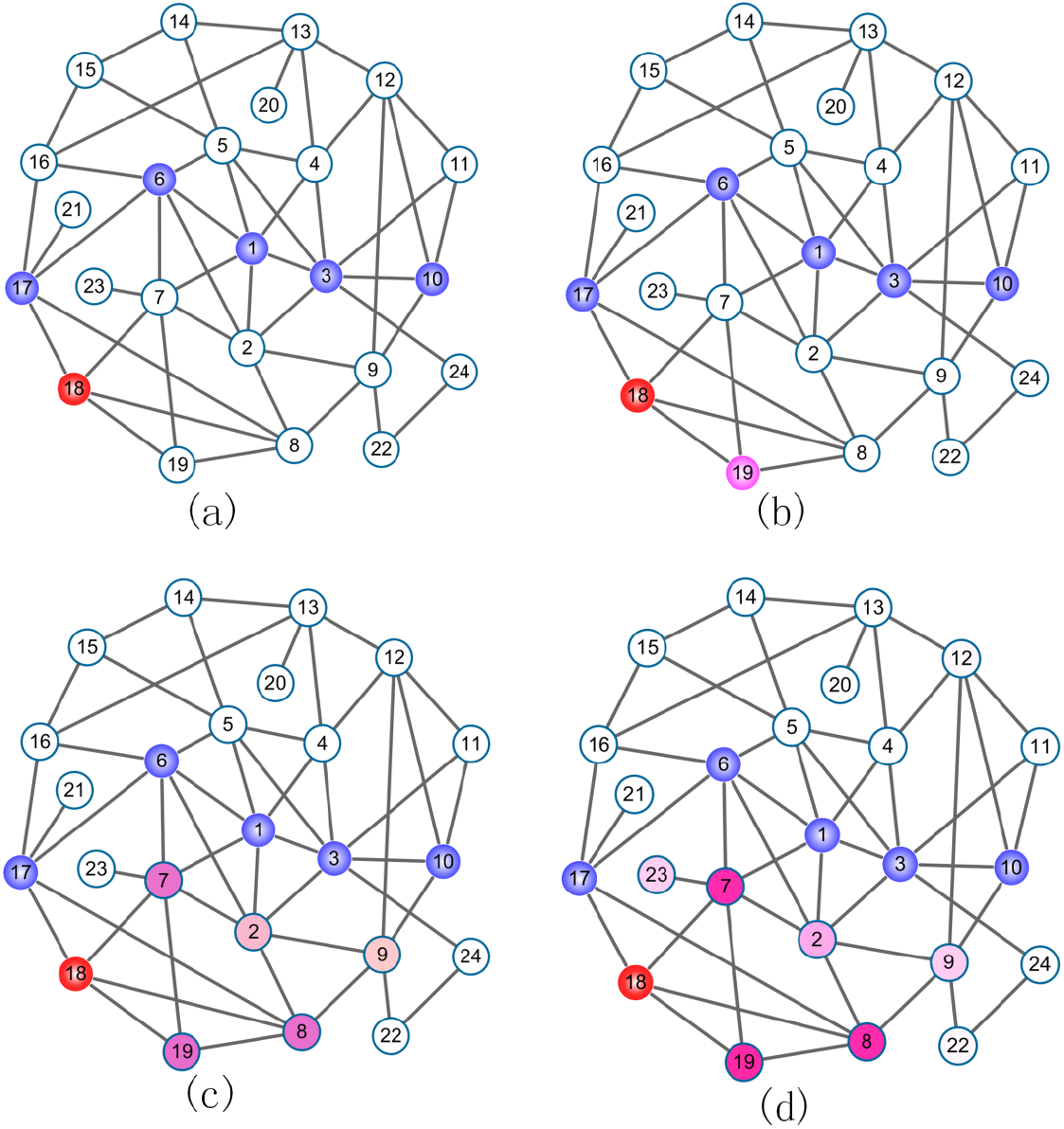
(Color online)A toy network with 24 nodes. (a) The snapshot includes 5 recovered nodes, i.e., 1, 3, 6, 10, 17, and 1 infected node, i.e., node 18, (b) a certain spreading simulation result from snapshot, only node 19 is infected when spreading achieves steady state, (c) average result on 10000 simulations from snapshot, and (d) result of probability prediction model from snapshot. In (c) and (d), the shades of nodes indicate the probability to be infected when spreading achieves steady state.

## Results and Discussion

To simulate the spreading process on networks, we employ the Susceptible-Infected-Removed (SIR) model [36]. In a network, we randomly select one node as the initial spreader. The information from this node will infect each of this node’s susceptible neighbors with probability *µ*, namely the infection probability. After infecting neighbors, the node will immediately become recovered (i.e., the recovering probability is 1). The new infected nodes in next step will infect their neighbors as the initial node. If it is not specially stated, we take the snapshot after five steps of spreading from the initial node as the known information.

We test our method on synthetic and real networks. Synthetic networks are Wattes-Strogatz (WS) networks [37], Barabási-Albert (BA) networks [38] and Given-Newman (GN) community networks [39]. Each synthetic network has 4000 nodes and each GN community network has 40 communities. We will discuss our model on three aspects: (1) the effect of infected probability *µ*, (2) the effect of structure of networks, and (3) the effect of stage of snapshot.

### The effect of infected probability

Fig. 2 shows the predictability *χ* and correlation *ρ* under different infected probability *µ* on WS, BA and GN networks. Generally, the predictability and correlation get larger with *µ* getting larger. For very large *µ*, e.g., *µ* = 0.3, the predictability and correlation approach to 1 since most of nodes will be infected. From Fig. 2, it can be seen that there exists a transition point, in detail, the transition point at *µ* = 0.15 for WS network and at *µ* = 0.1 for GN network. This can be explained as follows: the information almost do not diffusion if *µ* is small (*µ <* 0.15 for WS networks and *µ <* 0.1 for GN network), and the infected nodes are highly random for different simulations. It is noted that in BA network, it almost do not exist transition point. It can be explained as follows: since the heterogenous of its topological structure, regardless the location of initial infected node, the information will easily reach to the node with large degree, eventually, reach to other nodes. Interestingly, if *µ* is very small (e.g., *µ* = 0.02), the correlation is getting large in BA network, as shown in Fig. 2(a). Actually, for very small *µ*, just only a few snapshots have infected nodes, the results have no statistical significance. Besides, the distribution of correlation *ρ* under the results of 200 independent runs are listed in Figs. 2(d-f). From these three subfigures, it can be seen that the distributions of correlation *ρ* of BA and GN networks are similar, while that of WS network are generally large comparing with BA and GN networks.

**Fig 2.**
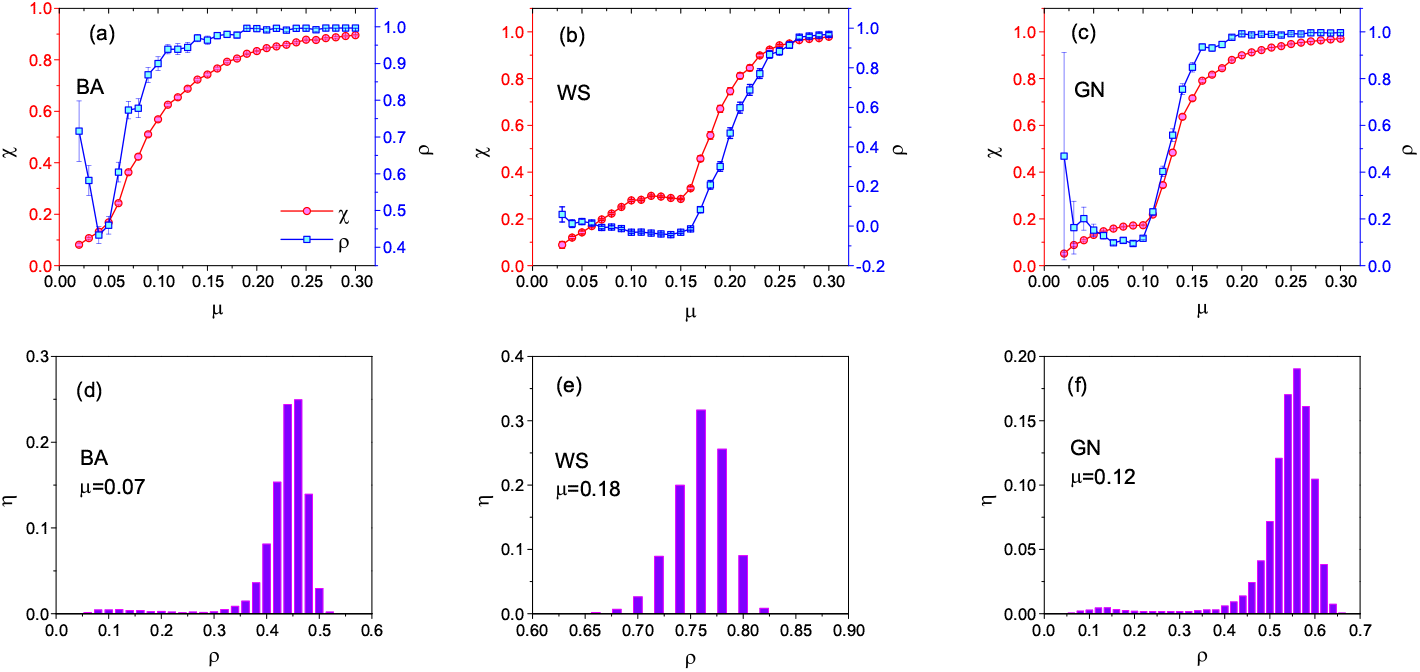
The predictability *χ* and correlation *ρ* under different infected probability *µ* on (a) BA, (b) WS and (c) GN networks. The distribution of Pearson coefficient in (d) BA, (e) WS and (f) GN are shown. The network parameters are *N* = 4000, ⟨*k*⟩ = 10, *p* = 0.1 for WS network, *N* = 4000, ⟨*k*⟩ = 10 for BA network, and *N* = 4000, ⟨*k*⟩ = 10, ⟨*k*_*in*_⟩ = 7 for GN network. The error bar in (a-c) and the distribution of correlation *ρ* in (d-f) are obtained by the results under 200 snapshots

### The effect of structure of networks

Fig. 3 shows the predictability and correlation for three types of networks with different structural parameters. For WS network, we study the effect of the rewiring parameter *p* on predictability and correlation. For BA network, we consider a variant of it in which each new node *u* connects to the existing node *v* with probability *P*_*u*_ = (*k*_*u*_ + *B*)/ Σ_*v*_ (*k*_*v*_ + *B*) [40, 41]. This modified model allows a selection of the exponent of the power-law scaling in the degree distribution *p*(*k*) ~ *k*^−*γ*^ with *γ* = 3 + *B/m* in the thermodynamic limit where *m* is the number of nodes should be connected when a new node is added and *B* is tunable parameter. With this network, we study the effect of *B* on predictability and correlation. For GN network, we study the effect of ⟨*k*_*in*_⟩ on predictability and correlation, where ⟨*k*_*in*_⟩ is the average internal degree of nodes in community. For a node *u* in community *C*, its internal degree 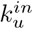 can de written as:

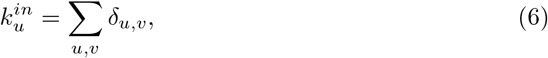

*δ*_*u,v*_ = 1 if *v* is also in community *C*, otherwise *δ*_*u,v*_ = 0. For standard BA network, i.e., *B* = 0, there are a few nodes with extremely large degree, the information can be spread out easily so long as it reaches to a node with large degree. So, it is relatively easy to predict which node will be infected in the future. As the *B* increases, the network evolves to random, a node getting infected or not will be hard to predict relatively, so the predictability and correlation decrease when *B* increases, as shown in Fig. 3(a). If rewiring probability *p <* 0.2, the information is hard to diffusion to other nodes since the WS network is almost regular, so it is hard to predict the infected nodes. When rewiring probability *p >* 0.2, the network has relatively strong random, the information reaches to other nodes easily, consequently, it is easy to predict the infected nodes, as shown in Fig. 3(b). In GN network, if average internal degree ⟨*k*_*in*_⟩ is larger, the community structure is clearer, correspondingly, the information is hard to escape the community boundary, and the predictability and correlation will getting worse, as shown in Fig. 3(c).

**Fig 3.**
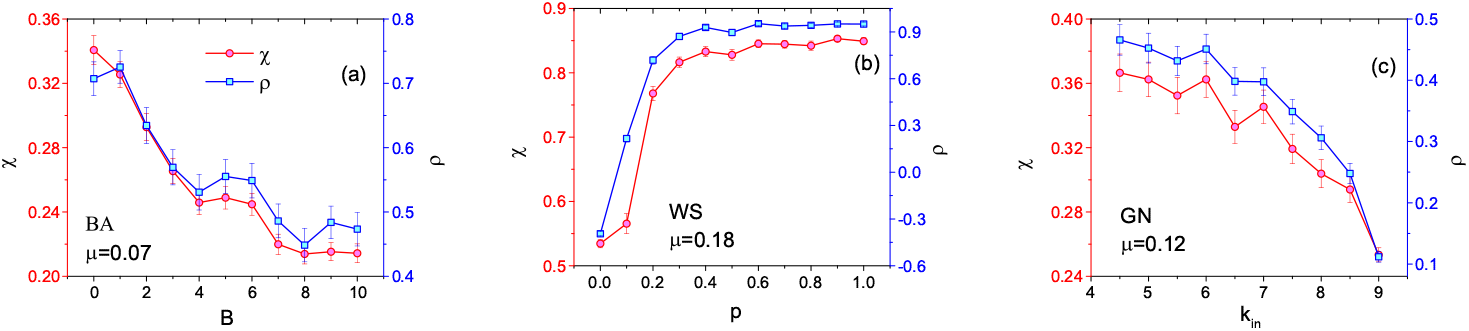
The predictability *χ* and correlation *ρ* for three types of networks with different structural parameters. In (a), B is a tunable parameter while generating network, (b) *p* is rewiring probability, and (c) ⟨*k*_*in*_⟩ is average internal degree.

Besides the network parameter listed above, the density of network, i.e., average node degree ⟨*k*⟩, also affects the predictability and correlation, as shown in Fig. 4. From Fig. 4, it can be seen that the *predictability* and correlation are small for small average node degree ⟨*k*⟩. Especially in WS and GN networks, for a large scope of average node degree (⟨*k*⟩ < 12 in WS and ⟨*k*⟩ < 8 in GN), the predictability and correlation are extremely small, there exists an obvious transition points, as shown in Fig. 4(a) and (c).

**Fig 4.**
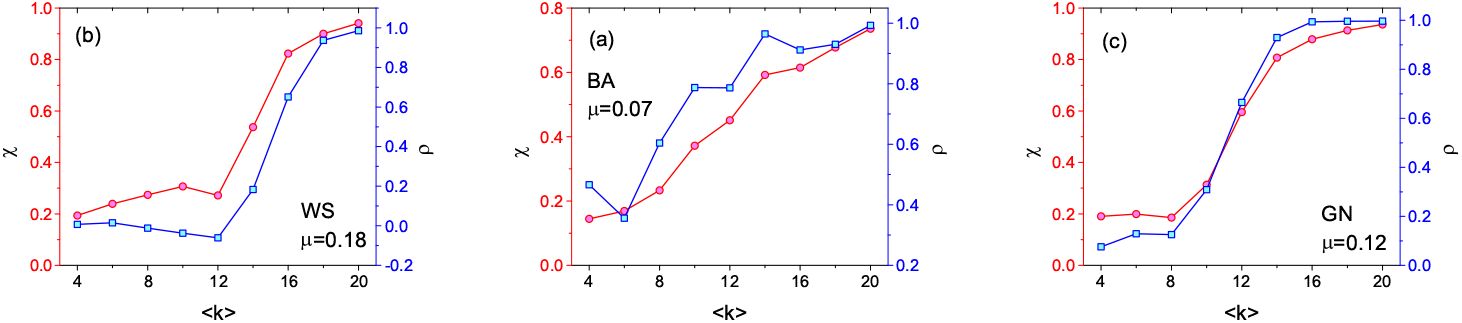
The effect of average node degree ⟨*k*⟩ on predictability *χ* and correlation *ρ*.

### The effect of stage of snapshot

We further analyze the predictability *χ* and correlation *ρ* under different stage of snapshot, as shown in Fig. 5. In Fig. 5, *T* is the spreading time of snapshot. Generally, it is difficult to estimate the infected rate precisely if just the snapshot in the early stage is given since there is little usable information, so, it is hard to predict the infected nodes. As *T* increases, more information could be used, the predictability *χ* and correlation *ρ* are getting better. In the late stage, many nodes of snapshot are infected or recovered, the left nodes are hard to be infected in reality, so the predictability *χ* and correlation *ρ* are getting worse, especially in BA network since most of all nodes are recovered.

**Fig 5.**
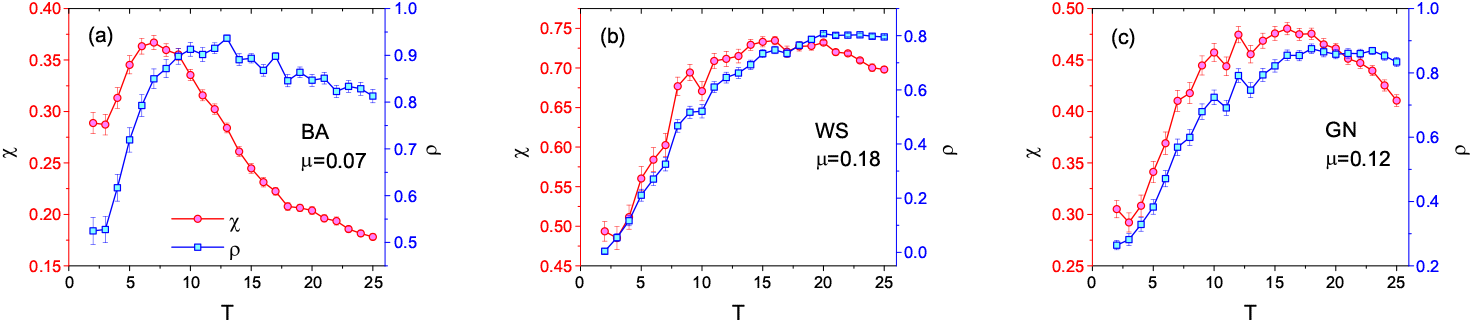
The predictability *χ* and correlation *ρ* under different stage of snapshot. Smaller *T* indicates earlier stage and larger *T* indicates latter stage.

Actually, if we analyze the number of infected nodes of snapshot in Figs. 2–5, we can find an interesting phenomena, as shown in Fig. 6. There is an obviously positive correlation between the number of infected nodes of snapshot and predictability *χ*. At the same time, it can be seen that the WS network has the strongest positive correlation while BA network has the weakest positive correlation under same number of infected nodes of snapshot. This might be universal, more infected nodes exist in snapshot, the information will be diffused easier, and so, it is more easy to predict the infected nodes in the future, correspondingly, the predictability *χ* will getting better.

**Fig 6.**
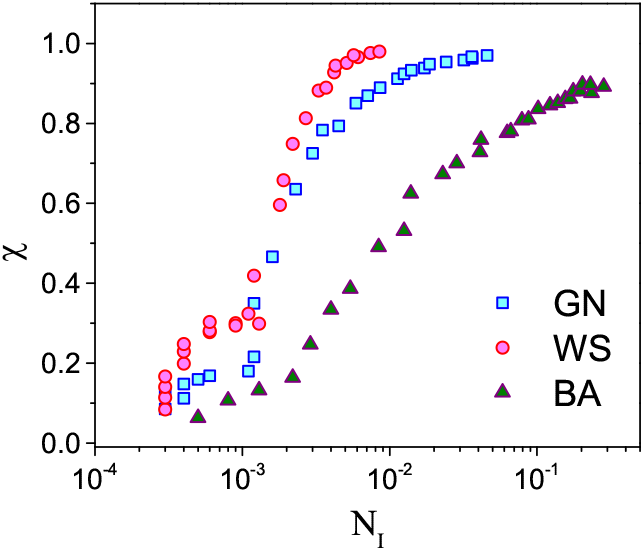
The correlation between the number of infected nodes *N*_*I*_ and the predictability *χ*.

Besides synthetic networks, we also analyze the predictability *χ* and correlation *ρ* on 11 real networks. The properties and analysis results on these real networks are shown in Table 1. From Table 1, it can be seen that the results are rather good, especial for the case of large *N*_*I*_, this is consistent with the results in Fig. 6. For networks Y2H and power, the predictability *χ* and correlation *ρ* are extremely low since *N*_*I*_ is very small. Actually, in these cases, there are few infected nodes in snapshot of spreading. Furthermore, the networks are very sparse, so, it is hard to predict the nodes being infected from snapshot in the future.

**Table 1.** The properties and analysis results on 11 real networks. The infected probability *µ* = 0.15.

| Networks | #Nodes | #Edges | $\chi$ | $\rho$ | $N_I$ |
| --- | --- | --- | --- | --- | --- |
| cond-mat | 39577 | 175693 | 0.7612 | 0.9430 | 0.0152 |
| astro-Ph | 16046 | 121251 | 0.8358 | 0.9426 | 0.0575 |
| email | 1133 | 5451 | 0.7854 | 0.9860 | 0.0628 |
| c.elegans | 453 | 2025 | 0.5577 | 0.9900 | 0.1143 |
| ecoli | 230 | 695 | 0.6110 | 0.9558 | 0.0509 |
| internet | 22963 | 48436 | 0.4525 | 0.9541 | 0.0625 |
| PGP | 10680 | 24316 | 0.6292 | 0.8074 | 0.0069 |
| TAP | 1373 | 6833 | 0.5998 | 0.5897 | 0.0101 |
| HEP | 7610 | 15751 | 0.4396 | 0.5975 | 0.0016 |
| Y2H | 1846 | 2203 | 0.2618 | 0.3214 | 0.0016 |
| power | 4941 | 6594 | 0.2375 | 0.2762 | 0.0003 |

## Conclusion

Up to now, most of researches mainly focus on the infection scale or threshold when they study the spreading dynamics in complex networks. However, following questions may be more important and interesting: Which nodes will be infected in the future and how to predict these nodes precisely? In this paper, we focused on this topic and presented a probability based prediction model to predict the infection nodes. Three synthetic and eleven real networks are used to evaluate the proposed model. Experimental results demonstrate that the model proposed could predict the infection nodes precisely in the sense of probability. In this paper, we just discuss the prediction model on static networks. The analyzing will get more difficult if the networks are evolving [42–44]. Furthermore, we analyze the effect of structure of networks, but we don’t consider the moving or self-protecting of individuals while disease outbreaks. Actually, as the diseases information makes individuals alert and take measures to prevent the diseases, the effective protection is more striking in small community [45]. We will study these more comprehensive cases deeply in the future.

## Acknowledgments

This work was partially supported by the National Natural Science Foundation of China with Grant No 61673085, by the Science Strength Promotion Programme of UESTC under Grant No. Y03111023901014006 and by the National Key Research and Development Program of China under Grant No. 2018YFB2100100.

